# Traveling waves support rhythmic attentional search

**DOI:** 10.64898/2026.03.16.712044

**Authors:** Yue Kong, Kirsten Petras, Philippe Marque, Rufin VanRullen, David M. Alexander, Laura Dugué

## Abstract

Brain activity unfolds across space and time, giving rise to spatiotemporal dynamics that can manifest as cortical traveling waves –smooth phase shifts propagating across the cortex. Although traveling waves have been observed across species and measurement scales, their functional role in human cognition remains largely inferred from correlational evidence. In healthy humans, large-scale neural dynamics must be studied non-invasively, which provides limited access to their neural origins. Here, we tested whether global traveling waves causally contribute to inter-areal communication during attention. Using transcranial magnetic stimulation (TMS), we induced long-range traveling waves detectable with electroencephalography and assessed their neural and behavioral consequences during an attentional search task. Double-pulse TMS applied over the right frontal eye field transiently disrupted ongoing global waves and selectively induced direction-and frequency-specific traveling waves propagating toward the occipital visual cortex. These TMS-induced anterior-to-posterior theta traveling waves were rhythmically modulated by stimulation latency relative to search-trial onset. Critically, behavioral performance exhibited the same rhythmic pattern, with improved attentional search performance when TMS-induced traveling waves were more prominent. Together, these findings provide causal evidence that large-scale traveling waves support inter-areal communication during rhythmic visual attention.

## Introduction

Traveling waves –smooth phase shifts across the cortex (Muller et al., 2018; Dugué & Chavane, 2025)– have been observed across species and scales and have been linked to cognitive processes (Zhang et al., 2018; Lozano-Soldevilla & VanRullen, 2019; Davis et al., 2020; Alamia et al., 2023; Fakche & Dugué, 2024; Fakche et al., 2024). In humans, electro-or magneto-encephalography (EEG, MEG) studies have reported wave-like patterns across the sensor array, but the significance of these observations is debated, partly due to uncertainties regarding their computational underpinnings (Grabot et al., 2025; Petras et al., 2025; Dugué & Chavane, 2025). This raises concerns about the functional relevance of these large-scale, non-invasively measured traveling waves. Here, we address this issue by using transcranial magnetic stimulation (TMS) to causally induce global traveling waves in EEG signals and examine their behavioral consequences.

Cortical traveling waves have been proposed as a canonical computation (Dugué & Chavane, 2025) that organizes neuronal processing across space and time in a variety of cognitive functions (for review Muller et al., 2018; Keitel et al., 2025), including attention (Alamia et al., 2023; Fakche et al., 2024). Attention itself is intrinsically spatiotemporal, and samples multiple objects in the sensory environment across space and time in a rhythmic manner, as reflected in theta-frequency (4-7 Hz) fluctuations in behavioral performance (VanRullen et al., 2007; Landau & Fries, 2012; Fiebelkorn et al., 2013; Senoussi et al., 2019; Galas et al., 2025; for review: Dugué & VanRullen, 2017; Fiebelkorn & Kastner, 2019; Kienitz et al., 2022). Such rhythmic modulations of performance can similarly be induced by time-locked perturbations such as with TMS. TMS over the right frontal eye field (rFEF), a key region of the attention network (Corbetta & Shulman, 2002; Corbetta et al., 2008; Scolari et al., 2015), has been shown to causally modulate oscillatory activity in visual perception and visuo-spatial attention tasks (Capotosto et al., 2009; Chanes et al., 2012, 2013; Stengel et al., 2021; for a comprehensive review see Vernet et al., 2014). When TMS is applied at different latencies relative to task events over the rFEF, visual attentional performance is periodically modulated over time (Dugué et al., 2019; Veniero et al., 2021). The rhythmic nature of visual attention aligns with neuroimaging evidence linking it to local, stationary neural oscillations in frontal (e.g., rFEF, Helfrich et al., 2018; Fiebelkorn et al., 2018) and occipital regions (Dugué et al., 2015; Landau et al., 2015; Plöchl et al., 2022). The involvement of both frontal and occipital regions is consistent with two-stage models of attention in which inter-areal iterative communication between low-level sensory processing regions and higher-order attention regions mediates attentional search (Palmer et al., 1993; Treisman, 1998; Itti & Koch, 2001; Deco et al., 2002). Yet, the neural mechanisms supporting such long-range coordination remain unclear.

Here, we investigate whether global traveling waves support inter-areal communication in visual attention. We tested the hypothesis that anterior-to-posterior global theta traveling waves have a causal, functional role in attentional search. We applied double-pulse TMS over rFEF (or over the vertex as a control site) at multiple stimulation latencies in an attentional search task while simultaneously recording EEG. This experiment’s behavioral data was previously used to demonstrate that rFEF-TMS rhythmically modulates attentional search performance as a function of stimulation latency (Dugué et al., 2019). Building on this established behavioral effect, we show here that TMS perturbed ongoing wave dynamics and selectively induced direction-and frequency-specific traveling waves. These TMS-induced traveling waves were themselves rhythmically modulated by TMS latency, and this neural modulation correlated with the previously reported rhythmic fluctuations in attentional search performance induced by TMS, thereby directly linking traveling wave dynamics to behavioral outcomes. Taken together, our results show that TMS-induced global traveling waves support inter-areal communication in rhythmic attentional search.

## Results

We tested the hypothesis that anterior-to-posterior global theta traveling waves have a causal functional role in attentional search. Participants performed an attentional search task with concurrent EEG recordings (Fig. 1a). Double-pulse TMS was applied over the rFEF at various latencies from search array onset. Stimulation of the vertex served as a control condition (Fig. 1b). We characterized global traveling wave activity in sensor space using a data-driven approach, complemented by time-frequency analyses and permutation-based statistical testing (Fig. 1c). We first assessed the prevalence of global traveling waves in TMS–EEG recordings, then examined how TMS transiently disrupted ongoing wave activity while inducing frequency-and direction-specific traveling waves. We then tested whether these TMS-induced waves were rhythmically modulated as a function of TMS latency. Finally, we compared this rhythmic modulation of TMS-induced traveling waves to rhythmic fluctuations in attentional search performance, such that more TMS-induced waves are correlated with improved behavioral outcome. Note that the behavioral effect observed in this experiment, i.e., rhythmic attentional performance, was previously reported in Dugué et al. (2019).

**Fig. 1.**
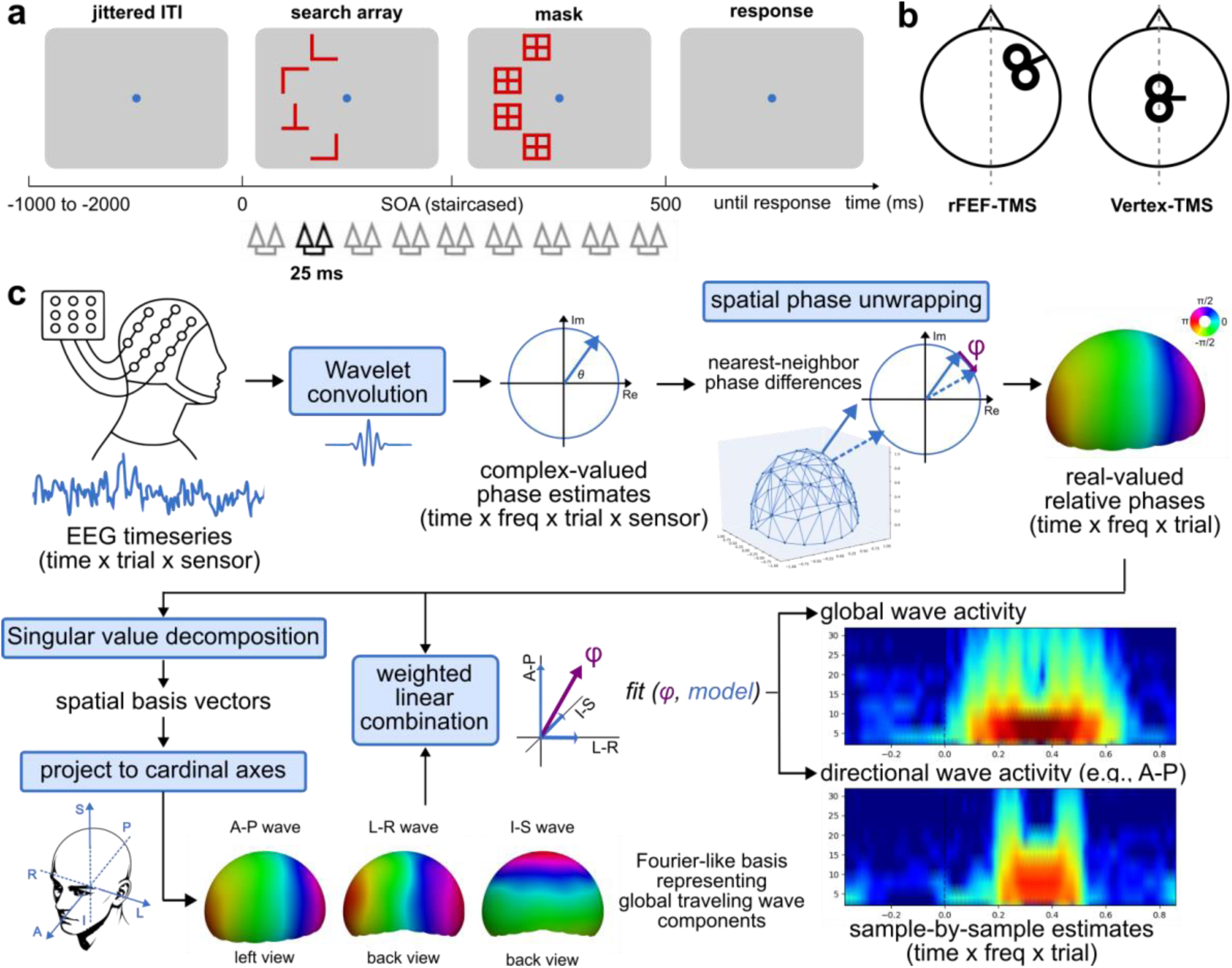
Experimental paradigm, TMS sites, and traveling wave analysis. **a** Attentional visual search task. Each trial began with a 1000-2000 ms jittered inter-trial interval (ITI). A four-item search array was briefly presented in the left visual field and masked after a staircase-determined stimulus-onset asynchrony (SOA), such that total visual stimulation lasted 500 ms. Participants reported the presence or absence of target “T” among distractor “L”s. A double-pulse TMS (25 ms inter-pulse interval) was delivered at one of nine delays during stimulus presentation. **b** TMS pulses were applied over the rFEF (left) or the vertex (right) as a control site. **c** Traveling wave analysis pipeline. Single-trial EEG data were wavelet-transformed to obtain complex-valued phase estimates, and then spatially unwrapped (via nearest-neighbor phase differences) into real-valued relative phases referenced to an arbitrary electrode (see Methods section). Singular value decomposition (SVD) was applied to the relative phases to extract spatial basis vectors. The first four spatial basis vectors were projected onto interpretable Cartesian dimensions of sensor space, yielding anterior-posterior (A-P), left-right (L-R), and inferior-superior (I-S) global traveling wave components. Relative phases were then reconstructed as a weighted linear combination of these components, providing sample-by-sample estimates of global or direction-specific wave activity across time and frequency (time-frequency wave maps).

### Theta-band power is selectively enhanced following TMS

To demonstrate the frequency specificity and the power of TMS-induced activity, we examined time-frequency representations and topographical distributions of induced power within trials in which TMS was applied at the latest latency (450 ms after search array onset). Grand-average power, computed across trials and sensors, revealed significant increases in the theta band (4.76-5.66 Hz) at two distinct post-stimulus intervals: an early window around 104-204 ms and a later window around 436-536 ms following search array onset (Fig. 2a and 2b). These windows were defined in a data-driven manner, independently of, and prior to the statistical comparisons described below, by identifying the peak theta power latency in each condition (rFEF-TMS and Vertex-TMS) using a 100ms-sliding-window average and centering a ±50ms-window on the corresponding peak.

**Fig. 2.**
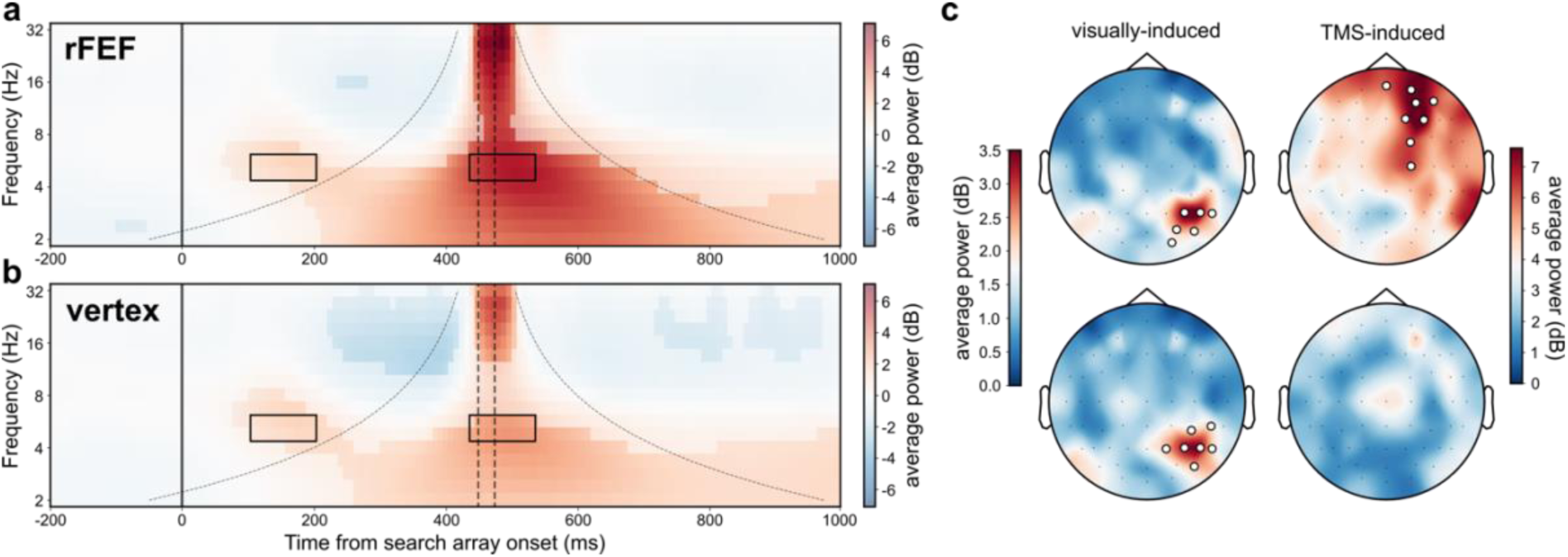
Theta-band power is selectively enhanced following TMS. **a** Grand-average time-frequency power (across trials and sensors) in the rFEF-TMS condition. The solid black line marks search array onset; dashed vertical lines indicate TMS single-pulse onsets. The cone of influence is overlayed (area potentially impacted by artifactual spread of the TMS-induced activity). Non-transparent regions denote significant clusters identified by permutation testing. Black boxes denote the early (104-204 ms) and late (436-536 ms) time-frequency windows of interest, defined in a data-driven manner from peak theta power latencies (see Methods). **b** Same as **a**, but for the Vertex-TMS (control) condition. **c** Scalp topographies averaged over the time-frequency windows outlined by black boxes in **a** (top row) and **b** (bottom row). Left column shows visually-induced activity averaged over 104-204 ms; right column shows TMS-induced activity averaged over 436-536 ms. Black circles outline electrodes within clusters of significant positive difference to zero (permutation testing).

To test whether the enhancement of theta power was selectively associated with TMS rather than a general effect of task timing, we conducted a direct cluster-based permutation test on the condition × time window interaction effect, defined as (rFEF-late − rFEF-early) − (Vertex-late − Vertex-early) (Maris & Oostenveld, 2007). This revealed a significant interaction (*p* = 0.0012), suggesting that the temporal profile of theta power differed reliably between TMS conditions across the scalp. A post-hoc simple-effect permutation test restricted to the late window (rFEF-late − Vertex-late) likewise revealed a significant result (cluster *p* < 0.001), indicating that the interaction was primarily driven by a condition difference in the late window. Condition-specific topographies within each window (Fig. 2c) were finally examined to visualize the underlying spatial patterns. The early and late time-frequency windows corresponded to visually-induced and TMS-induced activity, respectively, as illustrated by their corresponding scalp topographies (Fig. 2c). Theta power peaked over right occipital electrodes during the early window (104-204 ms) in both rFEF-TMS and Vertex-TMS conditions, consistent with visual stimulation in the left visual field. During the later window (436-536 ms), right frontal power peaked in the rFEF-TMS condition, whereas power increases in the Vertex-TMS condition were localized near electrode Cz and did not reach statistical significance (Fig. 2c; permutation tests, see Methods section).

Importantly, the visually-induced theta cluster fell outside the time-frequency cone of influence surrounding the TMS pulses confirming that it is not an artifact of the wavelet convolution. This cluster showed a clear temporal separation between visually-induced and TMS-induced activity; indicating the high temporal resolution of our time-frequency decomposition. Consistent with this interpretation, only the TMS-induced activity exhibited systematic temporal shifts across TMS latencies, whereas visually-induced theta power remained stable in both timing and magnitude across TMS latencies and stimulation conditions (Supplementary Fig. S1).

Together, our results suggest that TMS preferentially enhances theta-band activity, particularly following rFEF stimulation, providing a frequency-specific context for the subsequent observation of TMS-induced theta traveling waves. Consistent with this observation, the TMS-evoked potential over electrodes near the stimulation site showed sustained multi-cycle oscillatory activity at the individual-participant level (Supplementary Fig. S2a), which was masked in the grand-average TMS-evoked potential (Supplementary Fig. S2b), paralleling previous reports that averaging can obscure phase-organized single-trial activity (Alexander et al., 2013; Herring et al., 2015).

### Global traveling waves dominate EEG dynamics

Before testing our hypothesis that anterior-to-posterior global theta traveling waves have a causal functional role in attentional search, we first confirm that global traveling waves constitute a dominant component of EEG dynamics in the context of concurrent TMS-EEG experiments. To this end, global traveling wave components were empirically extracted for each participant using a singular-value-decomposition (SVD)-based approach (see Methods section). We define global traveling waves as phase dynamics with a spatial frequency of one cycle across the entire electrode array.

Across participants, the first four spatial basis vectors (main phase patterns; Fig. 1c) explaining the largest proportion of variance consistently exhibited a low spatial frequency, corresponding to approximately one cycle across the measurement array. Subsequent spatial basis vectors showed higher spatial frequencies and explained substantially less variance. By projecting these low-frequency spatial basis vectors onto three interpretable cardinal axes, we identified three global traveling wave components propagating along the anterior-posterior, inferior-superior, and left-right directions (Fig. 3a; see examples in Supplementary Fig. S3).

**Fig. 3.**
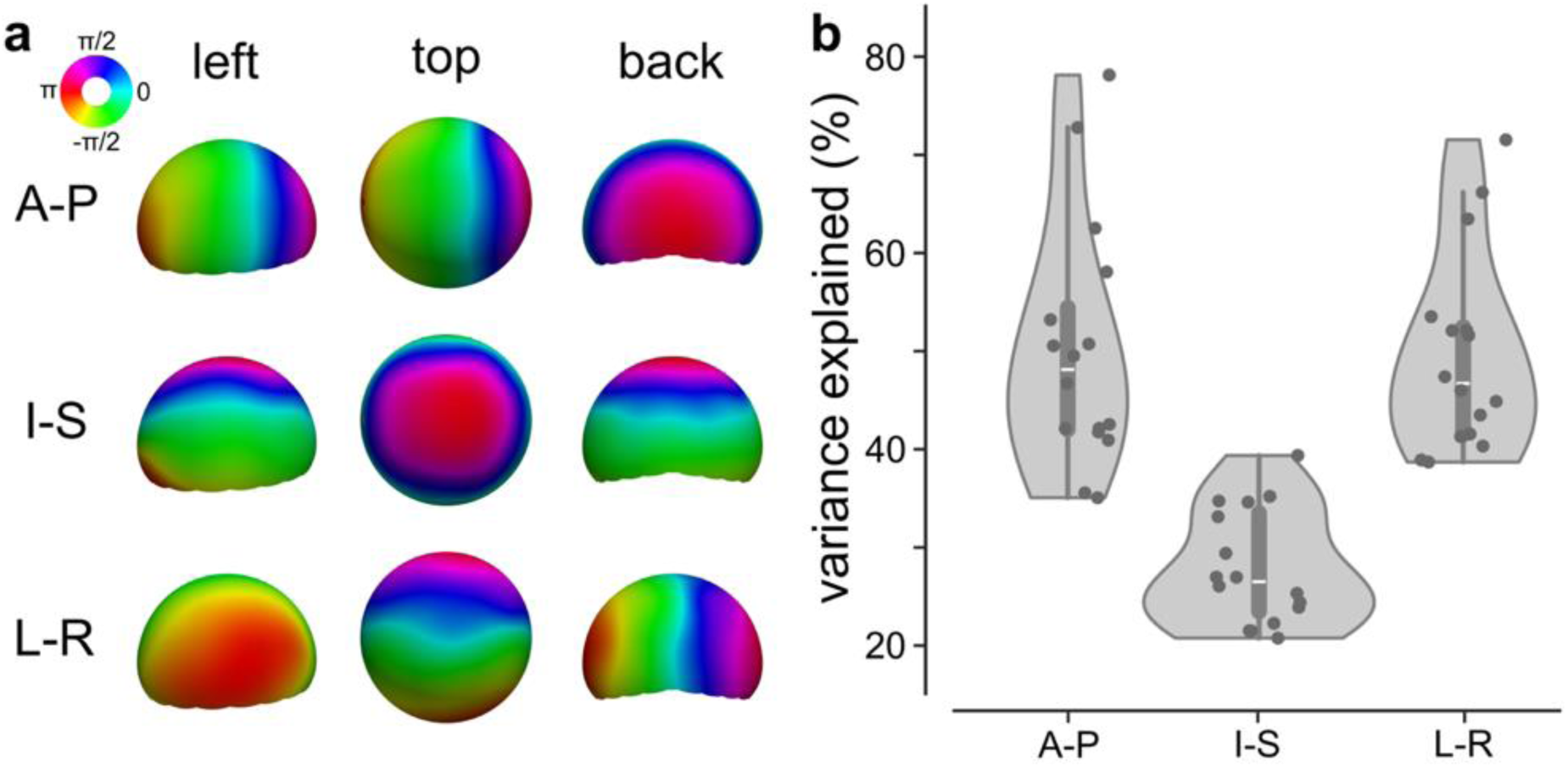
Global traveling waves dominate EEG dynamics. **a** Global traveling wave components projected onto cardinal axes (collapsed for participants). **b** Variance explained by each global traveling wave component after cardinal transformation across individual participants. A-P: anterior-posterior; I-S: inferior-superior; L-R: left-right.

The proportion of variance explained by each component is shown in Fig. 3b. The anterior-posterior component accounted for the largest share of variance, followed by the left-right component and the inferior-superior component. Low spatial-frequency global traveling waves account for the majority of EEG variance (58.13-69.76 %) and thus dominate cortical dynamics during concurrent TMS-EEG recordings. Notably, the summed variance explained by these three axes slightly exceeded 100 %, as the projection procedure renders the components marginally non-orthogonal.

### TMS transiently disrupts global wave activity

To assess the effect of TMS on ongoing traveling waves, trials were realigned to TMS onset, and global wave activity –quantified as the proportion of variance in the spatial phase pattern across electrodes explained by global traveling wave components (R^2^, therefore dimensionless; see Traveling Waves Analysis section)– was computed as a function of time and frequency. Permutation testing revealed a significant transient reduction in global wave activity time-locked to TMS onset in both the rFEF-TMS (Fig. 4a) and Vertex-TMS conditions (Fig. 4b). Here, global wave activity is a continuous variable averaged across trials, as demonstrated in the time-frequency plots at single-trial, single-participant, and grand-average levels (Supplementary Fig. S4). The grand-average significant cluster marks the time window where the TMS-induced disruption was most consistent across trials and participants, rather than reflecting an all-or-none event. A direct comparison between the rFEF-TMS and Vertex-TMS conditions using the same cluster-based permutation approach revealed no significant result (all cluster *p* > 0.05), indicating that the magnitude and time course of this disruption did not differ significantly between stimulation sites. In other words, the disruption by TMS reflects a general, non-specific consequence of the TMS pulse itself, largely independent of stimulation site. Additionally, this disruption was temporally specific to the TMS pulses, as evidenced by its systematic temporal shift across different TMS latencies (Supplementary Fig. S5), indicating that the observed effect reflects TMS-induced disruption of ongoing wave dynamics rather than task-related or spontaneous fluctuations.

**Fig. 4.**
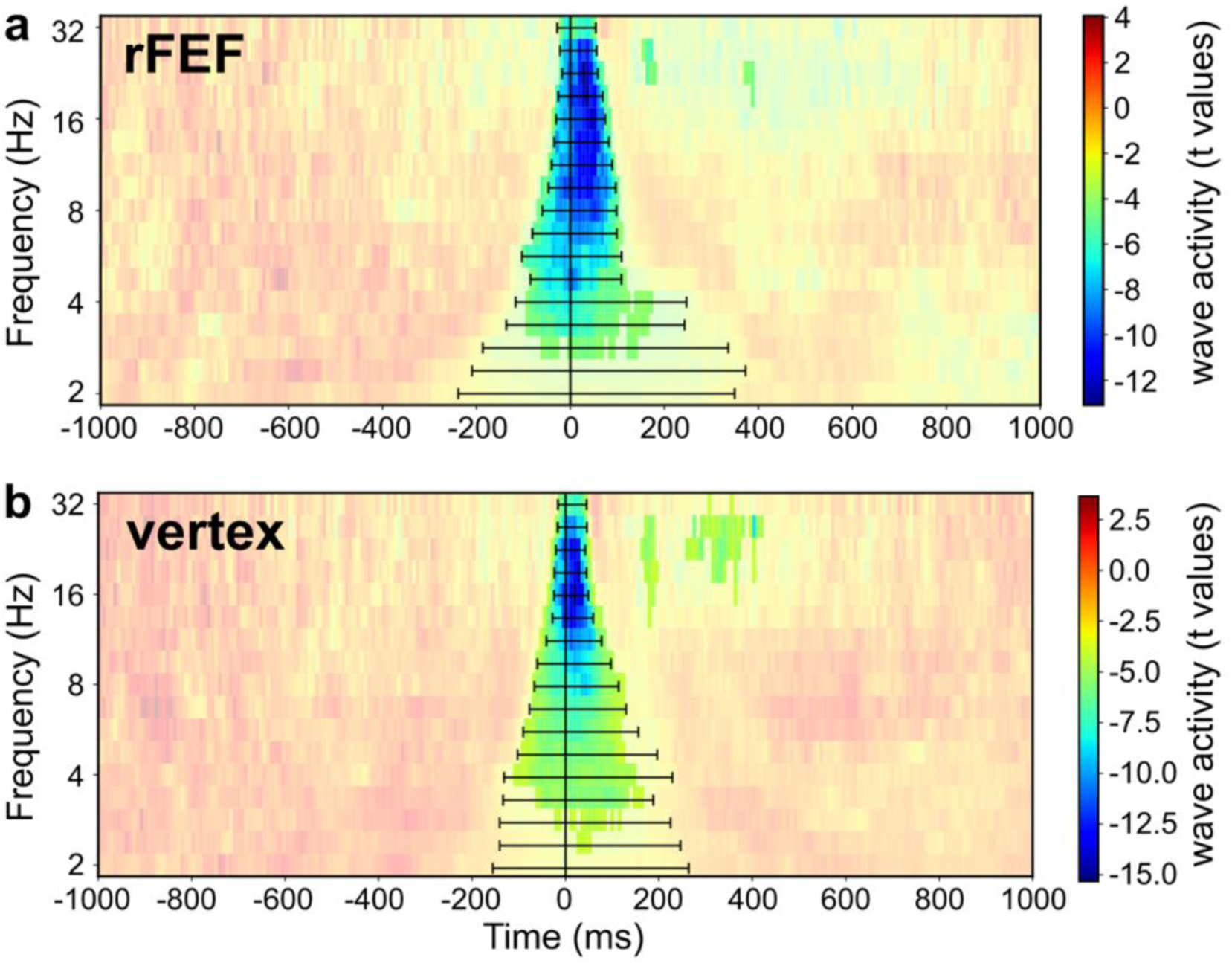
TMS transiently disrupts global wave activity. **a** Global traveling wave activity (t values) across time and frequency in the rFEF-TMS condition. The solid black line marks TMS onset (first pulse). Non-transparent regions indicate significant clusters identified by permutation testing. Horizontal error bars denote the full-width-at-half-maximum of disruption at each frequency, computed as the time window before and after the trough of maximal disruption, over which wave activity returned to half its maximal disrupted depth relative to baseline. **b** Same as **a**, but for the Vertex-TMS condition.

### rFEF-TMS selectively induces anterior-to-posterior global theta traveling waves

Based on our *a priori* hypothesis on anterior-to-posterior traveling waves, we specifically extracted traveling waves propagating from anterior to posterior direction across time and frequency. Despite the transient disruption of ongoing global traveling waves by TMS, stimulation of the rFEF led to a robust increase in anterior-to-posterior traveling wave activity in the theta frequency range (Fig. 5a; notice that this increase extends before TMS onset due to smearing from wavelet decomposition). In contrast, vertex stimulation produced a more confined increase in anterior-to-posterior wave activity that was largely restricted to the immediate post-TMS period (Fig. 5b). The statistical comparison between rFEF-TMS and Vertex-TMS was restricted *a priori* to the theta band, based on the previously established link between theta-band oscillations and rhythmic attentional sampling (e.g., VanRullen et al., 2007; Landau & Fries, 2012; Fiebelkorn et al., 2013; Senoussi et al., 2019; Galas et al., 2025; for review: Dugué & VanRullen, 2017; Fiebelkorn & Kastner, 2019; Kienitz et al., 2022), as well as on our own behavioral rhythms (previously reported in Dugué et al., 2019; data shown in Fig.6a) and data-driven identification of theta-band power peaks (Fig. 2a and 2b) prior to this analysis. This comparison revealed a significant cluster of enhanced anterior-to-posterior global theta traveling wave activity following rFEF stimulation, centered in the theta band (4.0– 5.7 Hz) and spanning approximately 60–220 ms after stimulation (Fig. 5c; cluster-based permutation test, *p* < 0.05). Note that the wave activity was estimated at each time sample (using short temporal window, two-cycle Morlet wavelets), thus not requiring wave persistence across multiple full oscillatory cycles. This allows that a significant cluster can be shorter than one full oscillatory cycle; the observation is that trial-averaged spatial phase organization reached significance; it does not mean that the difference is due to the presence of a single wavefront but rather the probability or strength of a traveling wave (Alexander et al., 2013; Alexander & Dugué, 2026). Here, single-trial anterior-to-posterior wave activity around TMS onset formed a continuous unimodal distribution in both the rFEF-TMS and Vertex-TMS conditions, with values concentrated at the lower end and with a tail toward higher values, as opposed to a bimodal, present/absent pattern (Supplementary Fig. S6).

**Fig. 5.**
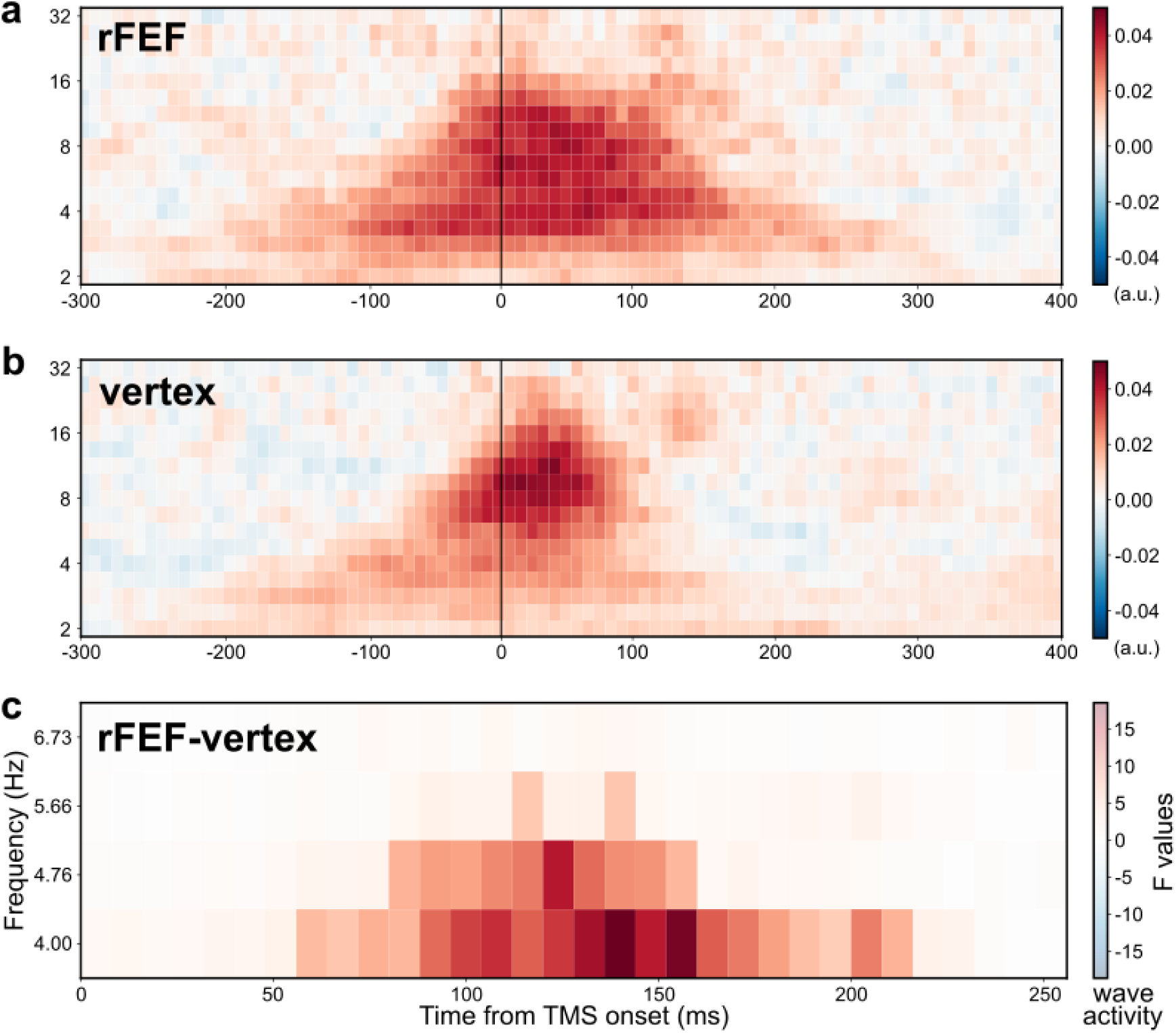
rFEF-TMS selectively induces anterior-to-posterior global theta traveling waves. **a** Anterior-to-posterior traveling wave activity across time and frequency in the rFEF-TMS condition. The solid black line marks TMS onset (first pulse). **b** Same as **a**, but for the Vertex-TMS condition. **c** Differences in anterior-to-posterior traveling wave activity between rFEF-TMS and Vertex-TMS conditions, expressed in F values. Non-transparent regions indicate significant clusters identified by permutation testing.

This effect was specific to the anterior-to-posterior axis, as analogous analyses along other directions (posterior-to-anterior, and both directions of the left-right and inferior-superior axes) did not reveal significant effects (see Supplementary Figs. S7 and S8). In comparison, the same analysis restricted to the alpha band, and often linked to perception and attention (Worden et al., 2000; Thut et al., 2006; Händel et al., 2011; Senoussi et al., 2026), did not show any significant difference between rFEF-TMS and Vertex-TMS condition. Anterior-to-posterior and posterior-to-anterior theta activity was finally examined in the pre-TMS window (−1000 to −250 ms relative to stimulation onset, avoiding overlap with the Wavelet cone of influence). A 2×2 repeated-measures ANOVA (Wave direction: anterior-to-posterior and posterior-to-anterior; TMS Condition: rFEF-TMS and Vertex-TMS) on theta wave activity within this window revealed a significant main effect of Wave direction (*F*(1,15) = 656.757, *p* < 0.0001), while neither the main effect of TMS Condition (*F*(1,15) = 0.973, *p* = 0.340) nor the Wave direction × TMS condition interaction (*F*(1,15) = 2.169, *p* = 0.162) reached significance. Post-hoc paired comparisons confirmed that anterior-to-posterior activity significantly exceeded posterior-to-anterior activity in both the rFEF-TMS and Vertex-TMS conditions (both corrected *p* < 0.0001). We argue that this result is coherent with the nature of the task, i.e., participant, throughout the task, must generally sustain their attention on the task at hand, coherent with top-down signals. Importantly, although both TMS conditions show pre-TMS anterior-to-posterior theta wave activity, rFEF-TMS selectively induces more anterior-to-posterior theta activity than Vertex-TMS within the post-stimulation window.

Taken together, these results demonstrate that TMS applied over the rFEF (compared to the vertex) selectively induces theta-band traveling waves propagating along the anterior-to-posterior direction, presumably from the rFEF to occipital visual regions, beyond nonspecific effects of TMS on overall wave activity, and beyond non-TMS related wave activity.

### rFEF-TMS modulates theta traveling waves rhythmically at 6.7 Hz

The behavioral data from this experiment were previously analyzed and reported in Dugué et al. (2019). This work demonstrated that TMS over the rFEF rhythmically modulates visual attentional search performance (measured as dprime) as a function of stimulation latency. Consistent with two-stage models of attention involving iterative communication between visual areas and higher-order regions, we then assessed whether global traveling waves support inter-areal communication in rhythmic attentional search. Specifically, we examined anterior-to-posterior global theta traveling waves as a function of TMS latency, and correlated TMS-induced wave activity with behavioral outcomes to assess the causal functional role of TMS-induced theta traveling waves in attentional search.

Wave activity was first averaged within the previously identified significant cluster between 60-220 ms (Fig. 5c) for each TMS latency and condition. Two-way repeated-measure analysis of variance (ANOVA) showed a significant main effect of TMS latency (*F*(8) = 2.376, *p* = 0.021), a significant main effect of TMS condition (*F*(1) = 12.246, *p* = 0.003), and a significant interaction between TMS latency and condition (*F*(8) = 2.058, *p* = 0.045). This result indicates that anterior-to-posterior global theta wave activity varies as a function of TMS latency, and that this effect is different between stimulations of the rFEF and the vertex.

We further examined the wave activity differences (rFEF-TMS minus Vertex-TMS) as a function of TMS latency (Fig. 6a). Spectral analysis using Fast Fourier Transform revealed a significant peak at 6.7 Hz, indicating that TMS-induced anterior-to-posterior global theta traveling waves are rhythmically modulated at this frequency (Fig. 6b). This modulation falls within the theta band and matches the periodicity previously reported for TMS effects on behavioral performance (Fig. 6a; note that more noise is expected in early latencies due to masking effects). Critically, the 6.7 Hz modulation was in phase across participants (Fig. 6b; Rayleigh test for circular data showed significant non-uniform distribution: *Z* = 6.1, *p* = 0.001).

**Fig. 6.**
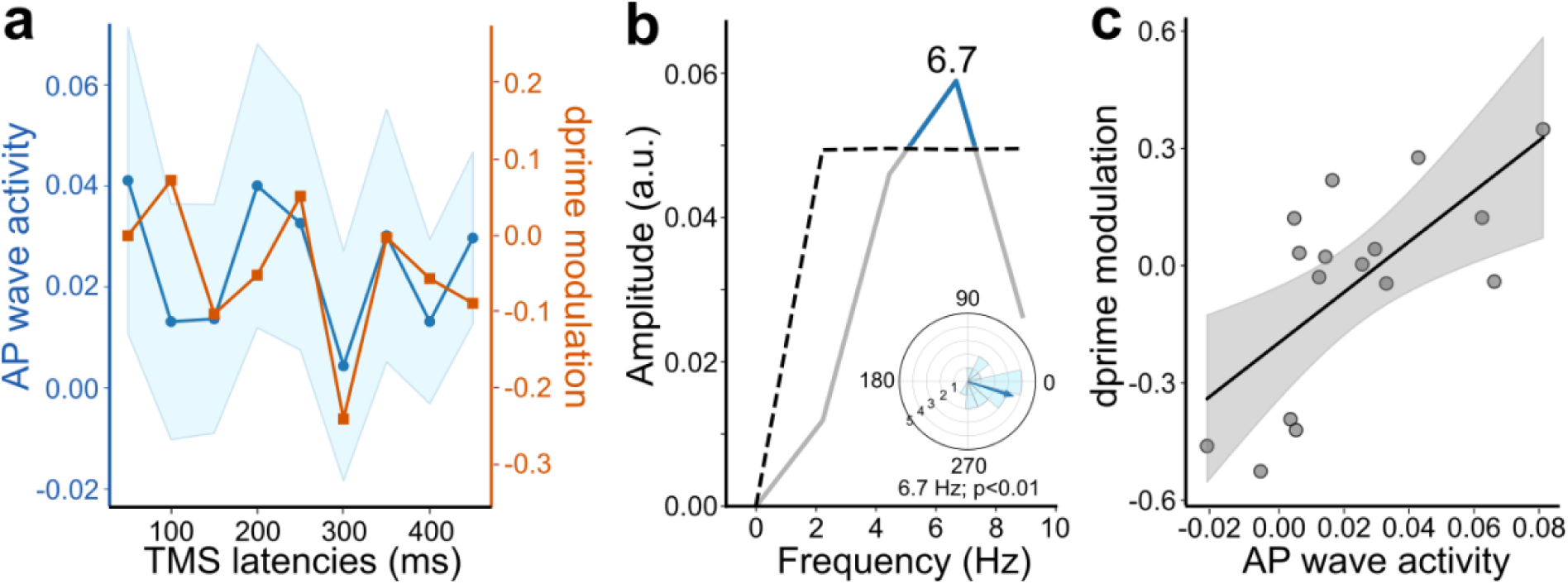
rFEF-TMS modulates theta traveling waves rhythmically at 6.7 Hz. **a** Both anterior-to-posterior (AP) wave activity (blue lines) and dprime modulation (orange lines) are represented as a function of TMS latencies from the search array onset. Shaded area indicates 95% confidence interval. **b** Amplitude spectrum obtained from Fast Fourier Transform of averaged AP wave activity. Dashed line marks statistical threshold (*p* = 0.05) from permutation test. Polar histogram of individual participant phases (light bins) and their average (blue arrow) at the peak frequency (6.7 Hz). **c** Correlation between AP wave activity and dprime modulation across participants. The solid line indicates the linear regression fit, with shaded area representing the 95% confidence interval. A significant positive correlation was observed (*r* = 0.677, *p* = 0.004).

Finally, we revealed a positive correlation between TMS-induced anterior-to-posterior global theta traveling waves and dprime modulation across individual participants (Fig. 6c; Pearson’s *r* = 0.677, *p* = 0.004). To assess whether this correlation was specifically attributable to traveling wave dynamics rather than co-occurring theta power modulations, we repeated this analysis using theta-band power, averaged within the same time-frequency window, in place of wave activity. Unlike wave activity, theta power showed no significant correlation with dprime (*r* = 0.321, *p* = 0.226). This dissociation indicates that the observed correlation between TMS-induced theta activity and behavior is specific to traveling wave dynamics rather than a general consequence of theta power modulation.

## Discussion

This study investigated the causal functional role of global traveling waves in inter-areal communication during visual attention. By applying double-pulse TMS over the rFEF at multiple latencies in an attentional search task while recording EEG, we show that TMS pulses transiently interrupt ongoing global wave dynamics and selectively induce direction-and frequency-specific traveling waves. Critically, rFEF stimulation elicits anterior-to-posterior global theta traveling waves that are rhythmically modulated as a function of TMS latency during the task, consistent with rhythmic fluctuations in attentional search performance.

The TMS-induced traveling waves reported here cannot be explained as a non-specific byproduct of the TMS pulse. On the one hand, the tested hypotheses are supported by carefully designed control conditions: 1) to test if double-pulse TMS over the rFEF selectively induced theta traveling waves, we used active stimulation at the vertex to control for potential auditory, somatosensory and muscular effects produced by TMS; 2) to test whether the induced theta wave activity is modulated by stimulation latency relative to search array onset, we used multiple TMS latencies within each stimulation condition as internal control, excluding any potential confounds caused by different stimulation sites. On the other hand, the transient disruption of global wave activity following TMS and its subsequent recovery did not differ between the rFEF-TMS and Vertex-TMS conditions and instead, scaled systematically with frequency. This indicates that the disruption itself reflects a generic physiological consequence of the TMS pulse, distinct from the direction-and frequency-specific anterior-to-posterior theta wave that constitutes our primary finding.

Cortical traveling waves have been linked to cognitive functions, including visual attention (for review Muller et al., 2018; Dugué & Chavane, 2025). A recent EEG study observed that during the allocation of spatial attention, alpha-band traveling waves propagate more from frontal to occipital electrodes than in the opposite direction along the same path (Alamia et al., 2023). Neural stimulation studies have demonstrated that external electrical perturbations can evoke or modulate propagating activity patterns in the cortex of different temporal frequencies (Massimini et al., 2005, 2007; Wischnewski et al., 2024; Lee et al., 2023; Wei et al., 2024; Campbell et al., 2025; Haigh et al., 2025). A recent transcranial alternating current stimulation study further demonstrated that electrically-induced anterior-to-posterior alpha traveling waves can modulate cognitive performance (visual attention task and memory task; Lee et al., 2026). Our results show that focal magnetic stimulation by TMS can causally induce large-scale traveling waves from a defined point of origin that predict rhythmic behavioral performance, providing direct evidence that traveling waves can have a causal functional role in rhythmic attentional sampling.

Importantly, the traveling waves observed here are unlikely to reflect purely local activity generated at the stimulation site. Indeed, authors have raised concerns regarding the interpretability of apparent traveling waves observed in non-invasive recordings (for review Dugué & Chavane, 2025). A recent study analyzed the spatial-frequency spectrum of gray-matter activity in intra-cranial EEG recordings (a signal not subject to volume conduction) and demonstrated that low spatial frequencies dominate cortical dynamics (Alexander & Dugué, 2024). This argues against explanations based exclusively on localized sources or standing waves (Orsher et al., 2023; Zhigalov & Jensen, 2023). Instead, these results support the presence of large-scale traveling waves measurable at the scalp level, consistent with the present interpretation of global traveling waves coordinating inter-areal communication in visual attention.

Here, TMS was applied over the rFEF, which constrained the point of origin of the induced anterior-to-posterior traveling waves to that cortical region. The rFEF has long been shown to play a critical role in the attention network (for review Corbetta & Shulman, 2002; Corbetta et al., 2008) and be causally linked to endogenous attention (Fernandez et al., 2023). Stimulation of rFEF not only affects behavioral performance in visual attention tasks but also alters neural excitability and evoked responses in visual cortex (Taylor et al., 2007; Capotosto et al., 2009; Chanes et al., 2012, 2013; Quentin et al., 2015; Dugué et al., 2019; Veniero et al., 2021; Stengel et al., 2021; for review see Vernet et al., 2014). This is consistent with other studies showing that the visual cortex is causally involved in such attentional tasks (Dugué et al., 2011; 2015). Although our analysis pipeline does not directly determine the ending site of the traveling waves, the anterior-to-posterior propagation observed here suggests that TMS-induced traveling waves propagate toward occipital visual areas. Future studies will test this hypothesis by probing the endpoint of waves, for example by combining the present TMS-EEG approach with a second TMS coil targeting the visual cortex.

A key open question concerns the anatomical pathways underlying the non-invasively observed global traveling waves. The anterior-to-posterior propagation from rFEF to occipital cortex is consistent with the attention network (Corbetta & Shulman, 2002; Corbetta et al., 2008; Scolari et al., 2015), but the present data do not allow us to determine whether these waves propagate via direct cortico-cortical connections through intermediate regions (e.g., parietal cortex) or emerge from subcortical relays. In particular, the thalamus (e.g., pulvinar nucleus) interacts with rFEF and is thought to coordinate functional interactions across visual areas (Saalmann et al., 2012; Halassa & Kastner, 2017). Computational modeling studies further suggest that thalamo-cortical circuits can generate emergent traveling waves and shape their spatio-temporal properties (Alamia & VanRullen, 2019; Bhattacharya et al., 2021; Schwenk & Alamia, 2025). Interestingly, fronto-parietal waves have been reported with invasive recordings in non-human primates during working memory and reward-guided behavior but at a different temporal frequency (beta: Bhattacharya et al., 2022; Zabeh et al., 2023). Is the different frequency due to the different tasks or is it reflecting the structural constraints on the network (here, different spatial scales and species; see Dugué & Chavane, 2025 for review). More generally, cortical traveling waves may be constrained by the structural connectome, whose topology shapes their propagation pathways, preferred directions, and frequencies (Koller et al., 2024). If propagation occurs through intermediate cortical regions, corresponding gradients of oscillatory power or phase shifts across the scalp would be expected. Future work using more spatially resolved approaches (likely in invasive recordings) as well as computational work (see Grabot et al., 2025) will be necessary to test these possibilities and directly characterize the underlying pathways of traveling waves.

Our findings are consistent with established theories of visual attention. Visual search has been postulated as a two-stage process involving low-level feature extraction and higher-order attentional selection, supported by iterative communication between visual areas and higher-order regions (Palmer et al., 1993; Treisman, 1998; Itti & Koch, 2001; Deco et al., 2002). Such exploratory behavior was more recently formalized in the rhythmic sampling theory of attention, which proposes that attention explores the sensory environment rhythmically (for review VanRullen, 2016; Dugué & VanRullen, 2017; Fiebelkorn & Kastner, 2019; Kienitz et al., 2022). Numerous studies have reported theta-band fluctuations in behavioral performance and neural oscillations during attention tasks (VanRullen et al., 2007; Landau & Fries, 2012; Fiebelkorn et al., 2013, 2018; Song et al., 2014; Chota et al., 2018; Helfrich et al., 2018; Senoussi et al., 2019; Michel et al., 2022; Galas et al., 2025). Our observation of functionally relevant traveling waves within the same frequency range provides strong empirical support that theta traveling waves are the underlying neural substrates of rhythmic attentional sampling.

Several influential theories have been proposed to explain the neural mechanisms underlying inter-areal communication, e.g., binding-by-synchrony (Singer & Gray, 1995), communication-through-coherence (Bastos et al., 2015), gating-by-inhibition (Jensen & Mazaheri, 2010), nested oscillations (Bonnefond et al., 2017). While these theories account for key temporal characteristics of oscillatory activity across brain regions, they generally conceptualize brain oscillations as stationary or locally synchronized processes. Instead, the traveling wave perspective considers temporal changes of global patterns of activity over areas of cortex, here large-scale cortex accessible via EEG. Our findings argue in favor of traveling waves supporting inter-areal communication, and are consistent with the latest proposal (Dugué & Chavane, 2025) that they constitute a canonical computation underlying diverse brain functions.

## Methods

All data were originally collected as part of a previously published study, for which the behavioral results along with a detailed description of the experimental protocol have been published by Dugué et al. (2019). Here, we analyzed the EEG data and describe only the relevant materials and methods.

### Participants

Twenty-three participants (7 women, age range: 24-38 years) with normal or corrected-to-normal vision participated in the experiment. Two did not complete the experiment due to discomfort during stimulation, and the EEG could not be recorded for five other participants for technical reasons. The EEG data from 16 participants were included in the analysis. All participants met the inclusion criteria for TMS and provided written informed consent prior to participation. The study was conducted in accordance with the Declaration of Helsinki and was approved by the local Ethics Committee (Sud-Ouest et Outre-Mer I; Protocol 2009-A01087-50).

### Visual Search Task

Participants performed an attentional visual search task (Fig. 1a; same procedure as in Dugué et al., 2011; Dugué et al., 2015). Visual stimuli were displayed on a screen (36.5 × 27 degree of visual angle, °VA) positioned 57 cm from the participant in a dark room. On each trial, four stimuli of identical size (1.5 × 1.5°VA) were presented at a fixed eccentricity (6°VA) in the left visual field. The stimulus array consisted either of four distractors (‘L’s; target absent trials) or of three distractors and one target randomly in one of the four stimulus locations (‘T’; target present trials). All items were randomly oriented (0, 90, 180, or 270° from upright).

After an individually determined stimulus-onset asynchrony (one-up-two-down staircase) to achieve approximately 70% correct performance (mean ± standard deviation: 85 ± 8 ms), the search array was masked, such that the total duration of visual stimulation was 500 ms. Participants were instructed to report the absence or presence of target ‘T’ by key press, with an emphasis on accuracy and no time pressure. The intertrial interval was jittered between 1000 and 2000 ms to minimize cumulative effects of stimulation. Participants completed 26 blocks of 72 trials each, including one practice block, one staircase block, four blocks without TMS, ten blocks with TMS applied over the rFEF (rFEF-TMS) and ten control blocks with TMS applied over the vertex (Vertex-TMS). The present study focuses the analyses on the rFEF-TMS and Vertex-TMS blocks.

### TMS procedure

TMS pulses were delivered using a Magstim Rapid^2^ stimulator (biphasic; Magstim Company, Whitland, UK) equipped with a 70-mm figure-of-eight coil. Stimulation was remotely triggered via Matlab R2014 from the stimulus presentation computer (Mathworks Inc., Natick, MA). Stimulation intensity was initially set at 50% of maximal output and individually adjusted to just below the threshold for facial and temporal muscle activation (mean ± standard error to the mean: 52 ± 2%). The coil was positioned tangentially to the scalp, with its handle oriented about 45° backward and laterally.

The stimulation target over the rFEF was defined based on averaged Talairach coordinates (x = 31, y = −2, z = 47). For 11 participants, coil positioning was guided by individual T1-weighted anatomical MRIs (3T Philips) using these coordinates. For the remaining participants, for whom anatomical MRI scans were not available, the stimulation site was defined as the barycenter (between electrodes F2 and FC4) of the rFEF locations identified in the MRI-guided participants. To control for nonspecific TMS effects (e.g., clicking noise and tapping sensation), TMS was also applied over the vertex as the control condition (block-designed, counterbalanced), which was individually defined as the location of electrode Cz (O’Shea et al., 2004). We chose active control-site stimulation over sham stimulation (e.g., tilted or placebo coil) because our hypothesis is about the specific link between rFEF and traveling waves in attentional search. A vertex stimulation reproduces the auditory, somatosensory and muscle activation at a comparable intensity and has minimal involvement in task-related processes (Jung et al., 2016).

Double-pulse TMS (inter-pulse interval: 25 ms) was delivered at one of nine possible latencies relative to search array onset (50-450 ms, in 50-ms increments). Double-pulse TMS was used instead of single-pulse stimulation because paired pulses have been shown to produce stronger and more reliable behavioral effects, particularly in visual search (Juan & Walsh, 2003; Kalla et al., 2008; Dugué et al., 2011, 2015; Yan et al., 2016). Here, the use of multiple latencies serves as an ideal internal control (see also Juan & Walsh, 2003; Dugué et al., 2011, 2015, 2016) to test the hypothesis that TMS-induced wave activity is modulated by the stimulation latency.

### EEG recording and pre-processing

EEG data were recorded using a TMS-compatible 64-channel EEG cap (standard antiCAP snap; 10–20 system; EasyCap GmbH, Herrsching, Germany). Electrodes Fpz and FCz served as ground and reference, respectively. Signals were sampled at 5000 Hz and acquired with an online low-pass filter at 1000 Hz.

EEG data were preprocessed using custom scripts in combination with the MNE-Python toolbox (Gramfort et al., 2013), following recommendations from TMS-EEG Signal Analyzer’s pipeline for TMS-EEG analysis (Rogasch et al., 2017). TMS-related artifacts in the raw EEG time series were removed (-1 to 15 ms relative to each pulse) and replaced using cubic interpolation based on five samples (1 ms) on each side of the interpolated window. Bad channels were identified by visual inspection and interpolated using MNE-Python’s spline interpolation. Data were then lowpass filtered at 330 Hz (finite impulse response, FIR; Hamming window; zero-phase; 82.5 Hz transition bandwidth; 53 dB stopband attenuation; passband edge 330 Hz; cutoff frequency 371.25 Hz; filter length 0.040 s) before being down-sampled to 1000 Hz to avoid aliasing.

Continuous data were epoched from-2 to 5 s relative to search array onset, followed by manual rejection of bad epochs. To remove large-amplitude artifacts (e.g., TMS-evoked muscle artifacts), independent component analysis (FastICA, 15 components) was performed on concatenated runs separately for rFEF-TMS and Vertex-TMS conditions. Cubic interpolation around TMS pulses was then applied again using the same parameters. After high-pass filtering at 0.1 Hz (FIR; Hamming window; minimum non-linear phase; 0.10 Hz transition bandwidth; 53 dB stopband attenuation), a second FastICA was performed to remove residual artifacts (e.g., blinks, eye movements, and muscle activity). Cubic interpolation around TMS pulses was subsequently applied once more.

Finally, EEG data were re-referenced to the common average. Channels TP9, TP10 and Iz were excluded due to their too-far distance from the remaining electrodes for the spatial phase unwrapping (see Traveling Waves Analysis section) leaving a total of 59 electrodes for the analysis. To equalize trial counts across conditions and participants, analyses were restricted to the first 400 trials for each of the rFEF-TMS and Vertex-TMS conditions.

### Time frequency decomposition

Time-frequency representations were computed using Morlet wavelet convolution. Phase and amplitude were estimated using two-cycle Morlet wavelets at 17 logarithmically spaced center frequencies ranging from 2 to 32 Hz. To reduce computational load, the resulting time-series of phase and power were down-sampled by a factor of eight resulting in a sampling rate of 125 Hz.

Grand-average power was first computed by averaging across all 59 electrodes and trials. Power values were then normalized into decibels relative to a pre-stimulus baseline (−1000 to −600 ms from search array onset). Significant clusters of induced power against baseline activity were identified using a cluster-based permutation test (Maris & Oostenveld, 2007; 5000 permutations, two-tailed), with a cluster-forming threshold of *p* = 0.001 and a cluster-level significance threshold of *p* = 0.05.

Topographic maps of induced power (decibel-normalized) were computed by averaging power within selected time-frequency windows defined from the grand-average data. Peak frequencies were identified in a data-driven manner, independently of, and prior to the following statistical comparisons, thereby avoiding circularity. Peak frequencies were identified from power spectra averaged across time, separately for each TMS condition. For each frequency, a 100-ms time window centered on the peak latency was selected (outlined in Fig. 2a and 2b).

To test whether theta power enhancement was selectively related to TMS, we performed a direct cluster-based permutation test on the condition × time window interaction effect. For each participant and electrode, we computed the interaction contrast [(rFEF-late − rFEF-early) – (Vertex-late − Vertex-early)] and ran a one-sample cluster-based permutation test (Maris & Oostenveld, 2007; 5000 permutations, two-tailed, electrode-adjacency-based clustering, cluster-forming threshold *p* = 0.05). Having established a significant interaction (Nieuwenhuis et al., 2011), we performed a post-hoc simple-effect test restricted to the late window (rFEF-late − Vertex-late), using the same cluster-based permutation procedure and threshold. Condition-specific topographies (Fig. 2c) were subsequently examined to characterize the spatial distribution of theta power increases relative to baseline within each condition and time window, using a separate cluster-based permutation test against baseline for each condition (one-tailed, testing for power increases relative to baseline; cluster-forming threshold *p* = 0.05).

### Traveling Wave Analysis

Cortical traveling waves were analyzed in single-trial EEG data using an SVD-based approach (Alexander et al., 2013). Complex-valued phase estimates (Φ) obtained from wavelet convolution were first converted, at each sample (time × frequency × trial × sensor), into real-valued relative phases (Ψ, relative to an arbitrary electrode) by computing nearest-neighbor phase differences across the electrode array and then reintegrating these differences in radians (spatial phase unwrapping; Fig. 1c). Specifically, for neighboring electrodes *i* and *j* (determined by Delaunay triangulation), their phase difference was defined as:

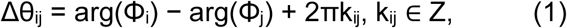

with integer k_ij_ chosen such that Δθ_ij_ ∈ [−π, π). The relative phase at each electrode was then reconstructed by cumulatively adding these nearest-neighbor differences (Δθ) according to:

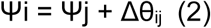

starting from an arbitrary reference electrode. These nearest-neighbor phase differences were sequentially integrated into an initially edgeless graph, in ascending order of |Δθ_ij_|. When the condition |Δθ_ij_| << π is not satisfied everywhere, inconsistencies may arise such that Ψi− Ψj = δΦij + 2kπ for some integer k ≠ 0. In data where phase varies smoothly across electrodes, the number of such inconsistencies is small relative to the total number of nearest neighbors. These inconsistencies may reflect either genuine phase discontinuities or measurement noise (Spagnolini, 1995). Integrating phase differences by increasing |Δθ_ij_| preferentially localizes such errors to regions exhibiting the largest phase differences, corresponding to legitimate discontinuities or elevated noise.

At the single-participant level, SVD was applied to the relative phase data to reduce dimensionality and extract dominant spatial patterns of phase dynamics across the scalp. For each participant, relative phase values were arranged into a matrix A ∈ R^Ne×Ns^, where N_e_ denotes the number of electrodes and N_s_ the number of samples collapsed across time, frequency, trial, and TMS condition. The matrix A was decomposed by SVD as

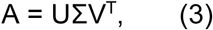

where U contains the left singular vectors (LSVs), Σ is a diagonal matrix of singular values σ_r_ ordered in descending magnitude, and V contains the right singular vectors. The LSVs represent orthogonal spatial basis vectors that capture dominant phase gradients across the electrode array, whereas the corresponding singular values (σ) quantify the square root of the variance explained by each vector. By definition, these empirically-derived vectors are orthogonal and can be treated as independent modes of the system; thus, weighted linear combinations of the LSVs can recover the original phase data without loss. Because standard Fourier methods cannot be easily applied to EEG’s irregular sensor layout and to the partially covered surface of a spheroid, so this SVD-based decomposition serves as an empirically-based Fourier analysis, expressing the phase pattern as a weighted sum of approximate spatial sinusoids of increasing spatial frequency (wave number k = 1, 2,…;Alexander & Dugué, 2026; Brunton & Kutz, 2019; Shinn, 2023). Unlike an analysis done on complex-valued phase (Alexander & Dugue, 2026), the real SVD on spatially unwrapped phases reduces this Fourier problem into one of weighted spatial gradients.

Spatial basis vectors were retained in descending order of explained variance, with the first three to four components typically corresponding to a spatial frequency of approximately one cycle across the electrode array (k = 1). This restriction to the lowest-order spatial component was deliberate: independent intracranial evidence indicates that cortical phase organization is dominated by low spatial frequencies across temporal bands including theta (Alexander & Dugué, 2026), well below the spatial scale associated with volume conduction and gyral folding. Because volume conduction acts as a spatial low-pass filter, this low-frequency-dominant regime is not created at the scalp, supporting the use of scalp EEG to detect genuine, non-artifactual long-wavelength traveling waves. These participant-specific spatial basis vectors represent global traveling wave components, each corresponding to an orthogonal direction of phase propagation across the scalp. Linear combinations of these spatial basis vectors reconstitute all possible global TW components.

To facilitate interpretation and comparison across participants, the individual spatial basis vectors were projected onto a common set of axes defined by electrode coordinates in three Cartesian dimensions (anterior-posterior, inferior-superior, and left-right; Fig. 3a). This procedure effectively rotates each individual’s spatial basis vectors to a shared cardinal reference frame, enabling direct cross-participant comparison. Prior to projection, phase values were normalized according to:

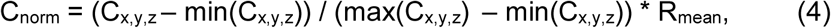

where C_x,y,z_ denotes the phase values expressed in electrode coordinate space and R_mean_ defines the mean range of the retained spatial basis vectors. Projection coefficients were then computed as

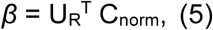

where U_R_ contains the first *R* spatial basis vectors obtained from SVD, with R determined by the number of global traveling wave components retained (here *R* = 4). Reconstructed phase dynamics were obtained as:

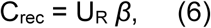

yielding sample-by-sample estimates of wave activity. We define wave activity as the proportion of variance in the measured spatial phase pattern explained by this reconstruction (R², bounded between 0 and 1), computed as the squared correlation between the measured phase vector and its reconstruction (C_rec_) at each time sample. This measure is computed independently at each individual time sample, after aggregation across all bases (global wave activity; Fig. 4) or, for direction-specific measures restricted to individual cardinal bases (e.g., anterior-to-posterior waves; Fig. 5).

Single-trial global wave activity was re-aligned to TMS onset and averaged across trials, followed by baseline correction using a pre-TMS interval (−1000 to −600 ms). For each condition, significant modulations of wave activity were assessed using two-tailed cluster-based permutation tests (Maris & Oostenveld, 2007; 5000 permutations), with a cluster-forming threshold of *p* = 0.001 and a cluster-level significance threshold of *p* = 0.05. The same cluster-based permutation approach was used to directly compare the rFEF-TMS and Vertex-TMS conditions.

TMS-related anterior-to-posterior wave activity was then compared between the rFEF-TMS and Vertex-TMS conditions using cluster-based permutation testing (Maris & Oostenveld, 2007; 5000 permutations, two-tailed, *p* = 0.05). Statistical testing was restricted to a pre-defined time-frequency window (4-7 Hz, 0-250 ms relative to TMS onset). The theta frequency band was selected *a priori* based on the previously established link between theta-band oscillations and rhythmic attentional sampling (VanRullen et al., 2007; Landau & Fries, 2012; Fiebelkorn et al., 2013; Senoussi et al., 2019; Galas et al., 2025; for review: Dugué & VanRullen, 2017; Fiebelkorn & Kastner, 2019; Kienitz et al., 2022), our own behavioral rhythms (data previously reported in Dugué et al., 2019), and the data-driven identification of theta-band power peaks in the grand-average power spectrum (Fig.2a and 2b); the temporal window was chosen based on previous time-frequency analyses showing maximal grand-average power in this interval (Fig. 2a and 2b). We also repeated the same cluster-based permutation analysis restricted to the alpha band to further strengthen the specificity of theta. To assess the directional distribution of theta wave activity independent of any TMS-induced effect, we additionally defined a pre-TMS window of −1000 to −250 ms relative to stimulation onset, selected to avoid overlap with the Wavelet cone of influence around TMS onset. Mean theta-band anterior-to-posterior and posterior-to-anterior wave activity within this window was tested in a 2×2 repeated-measures ANOVA with factors Wave direction (anterior-to-posterior vs. posterior-to-anterior) and TMS Condition (rFEF-TMS vs. Vertex-TMS), followed by post-hoc paired t-tests (Holm-corrected for multiple comparisons) to characterize simple effects.

To characterize the rhythmic modulation of TMS on anterior-to-posterior global theta traveling waves, wave activity was averaged within the significant cluster identified above, and differences between rFEF-TMS and Vertex-TMS conditions were computed as a function of TMS latency. Periodicity was assessed by applying a Fast Fourier Transform to the detrended time series (Kienitz et al., 2022). Statistical significance at each frequency was evaluated using a bootstrapping procedure. Specifically, TMS latency labels were randomly permuted within participants (100,000 iterations), and the Fast Fourier Transform was recomputed for each surrogate spectrum. For each frequency, we computed the 95^th^ percentile of the surrogate distribution obtained at the same frequency across all permutations. The empirical spectrum was considered significant at a given frequency if it exceeded this frequency-specific threshold (*p* < 0.05). For the frequency showing a significant amplitude peak, phase values were extracted for each participant to verify that the observed periodicity was in phase across participants, and the circular distribution of these phases was assessed using a Rayleigh test for non-uniformity. Finally, Pearson correlation coefficient was computed to assess the link between anterior-to-posterior waves and dprime modulation across participants. To assess whether it was specific to traveling wave dynamics rather than co-occurring theta power modulations, we repeated the same analysis using theta-band power, averaged within the same time-frequency window, in place of wave activity. It is worth noting that these analyses use orthogonal components of the signal, since temporal power is explicitly removed from the phase signal before TW analysis. Although wave activity was quantified at the single-trial level, behavioral performance was better summarized using dprime, a sensitivity measure free of decision biases (Green & Swets, 1966) that inherently requires trial aggregation. This metric isolates perceptual sensitivity from response bias and therefore constitutes a more reliable and theoretically appropriate measure for relating neural dynamics to behavior.

## Acknowledgements

This project has received funding from the European Research Council (ERC) under the European Union’s Horizon 2020 research and innovation programme (grant agreement No 852139 - Laura Dugué). We also thank Alexy-Assaf Beck for collecting the data for the original experiment (Dugué et al., 2019), as well as Andrea Alamia and Camille Fakche for their useful comments on the manuscript.

## Author contributions

L.D. and Y.K. designed the research. Y.K. and D.M.A. performed the analysis of the data. Y.K., D.M.A. and L.D. interpreted the data. Y.K. wrote the first version of the manuscript. All authors revised and edited subsequent versions of the manuscript.

## Data and code availability

All data and analysis code will be made publicly available upon acceptance of the manuscript. The MEG_Chicken toolbox (https://github.com/DugueLab/MEG_Chicken) was used by the first author as a training program for artifact detection (combined with training from the senior authors).

## Supplementary Information

**Supplementary Fig. S1.**
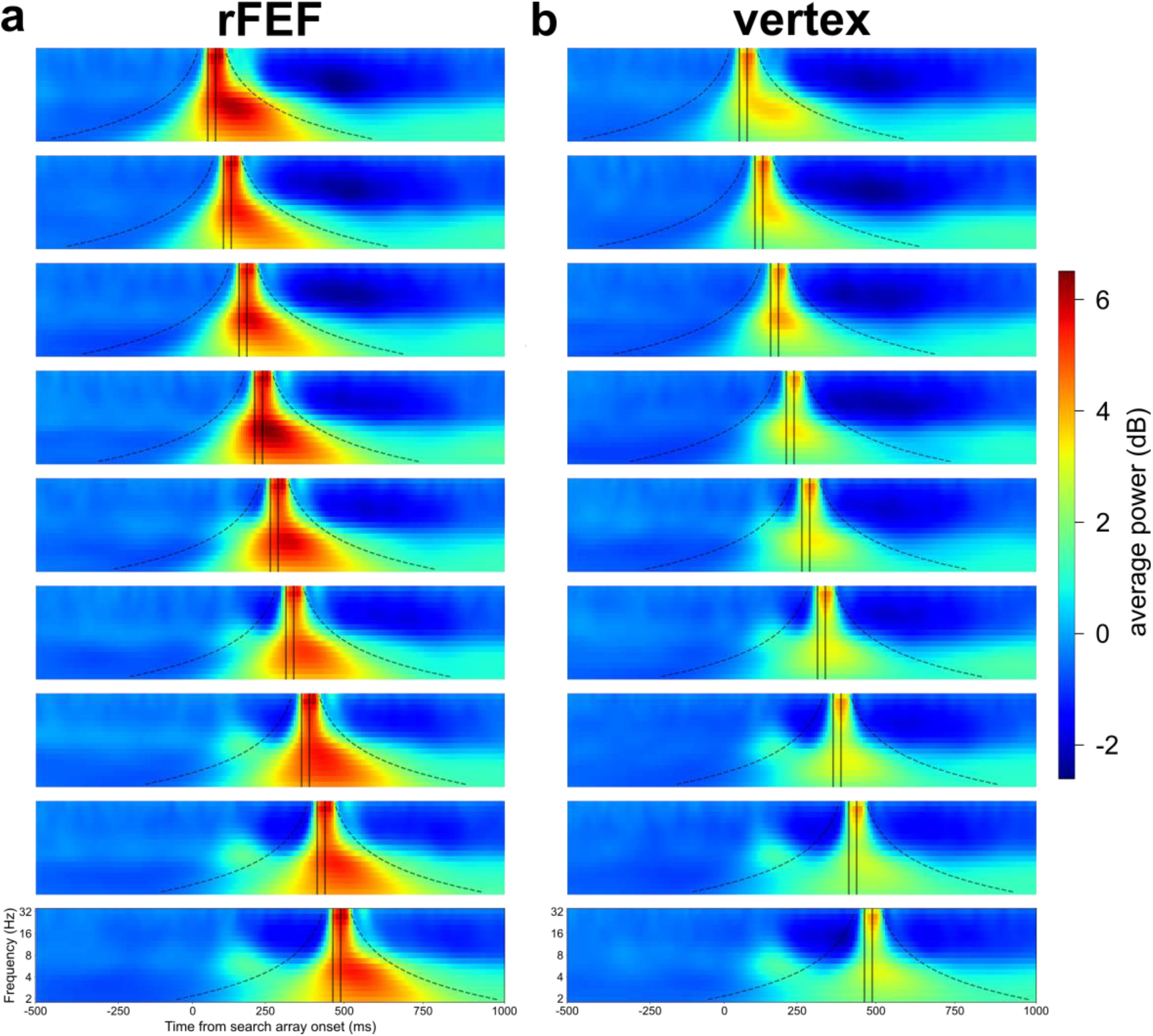
TMS-induced power systematically shifts across TMS latencies. **a** Grand-average time-frequency power (across participants, trials and sensors) in the rFEF-TMS condition. The solid black lines indicate single-pulse TMS onsets. The cone of influence is overlayed (area potentially impacted by artifactual spread of the TMS-induced activity). Only the TMS-induced activity exhibited systematic temporal shifts across TMS latencies, whereas visually-induced theta power (around 100-200ms) remained stable in both timing and magnitude across TMS latencies and stimulation conditions. **b** Same as **a**, but for the Vertex-TMS (control) condition.

**Supplementary Fig. S2.**
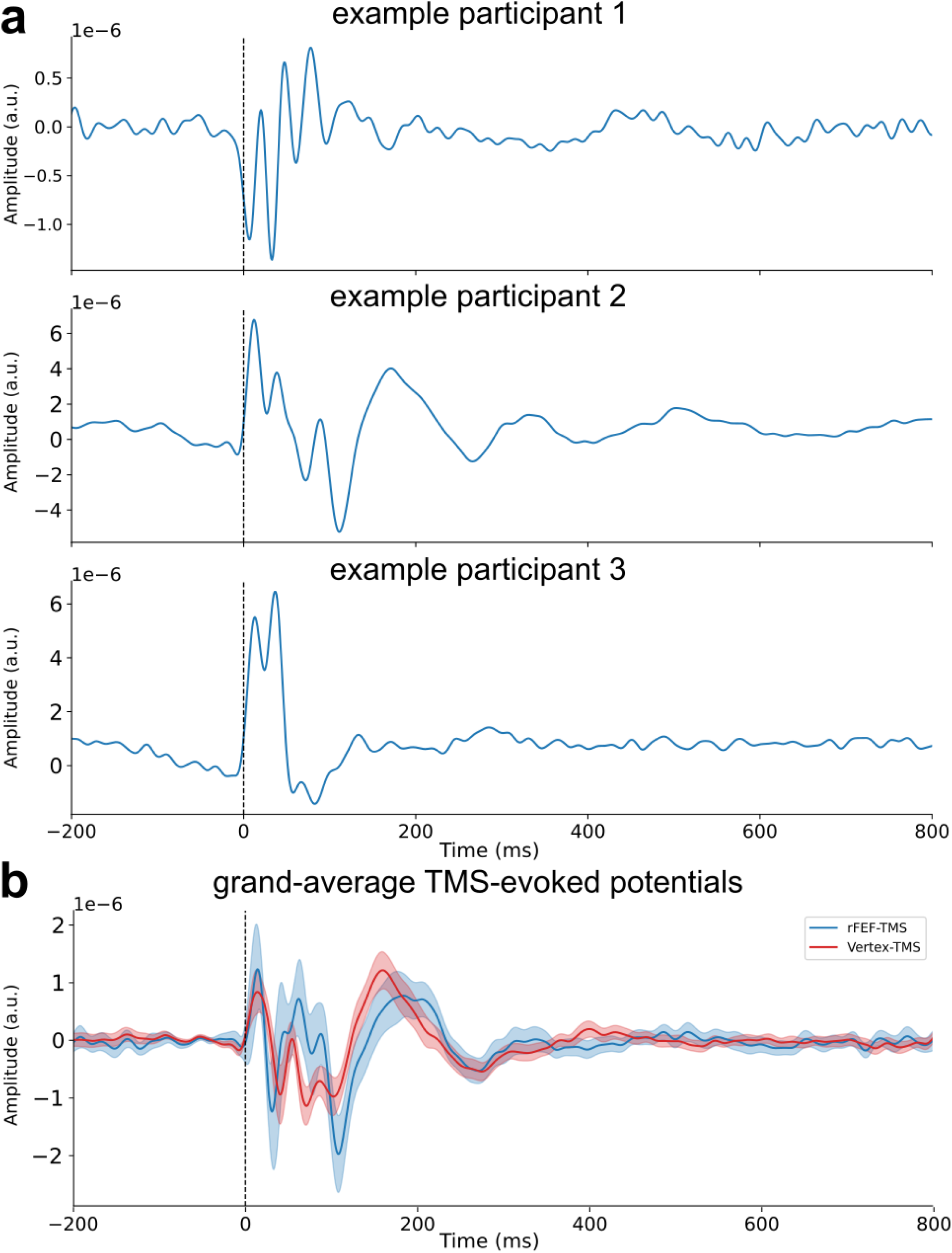
Individual TMS-evoked potentials reveal sustained oscillatory activity that attenuates in the grand average. **a** TMS-evoked potentials from three example participants (rFEF-TMS condition; band-pass filtered 3–45 Hz), time-locked to TMS onset (dashed line) and averaged over electrodes near the stimulation site. **b** Grand-average TMS-evoked potentials for the rFEF-TMS (blue) and Vertex-TMS (red) conditions. Shaded areas denote the standard error of the mean across participants. Individual-participant TMS-evoked potentials reveal sustained oscillatory activity, which is reduced in the grand average due to variability in oscillation onset and phase across participants.

**Supplementary Fig. S3.**
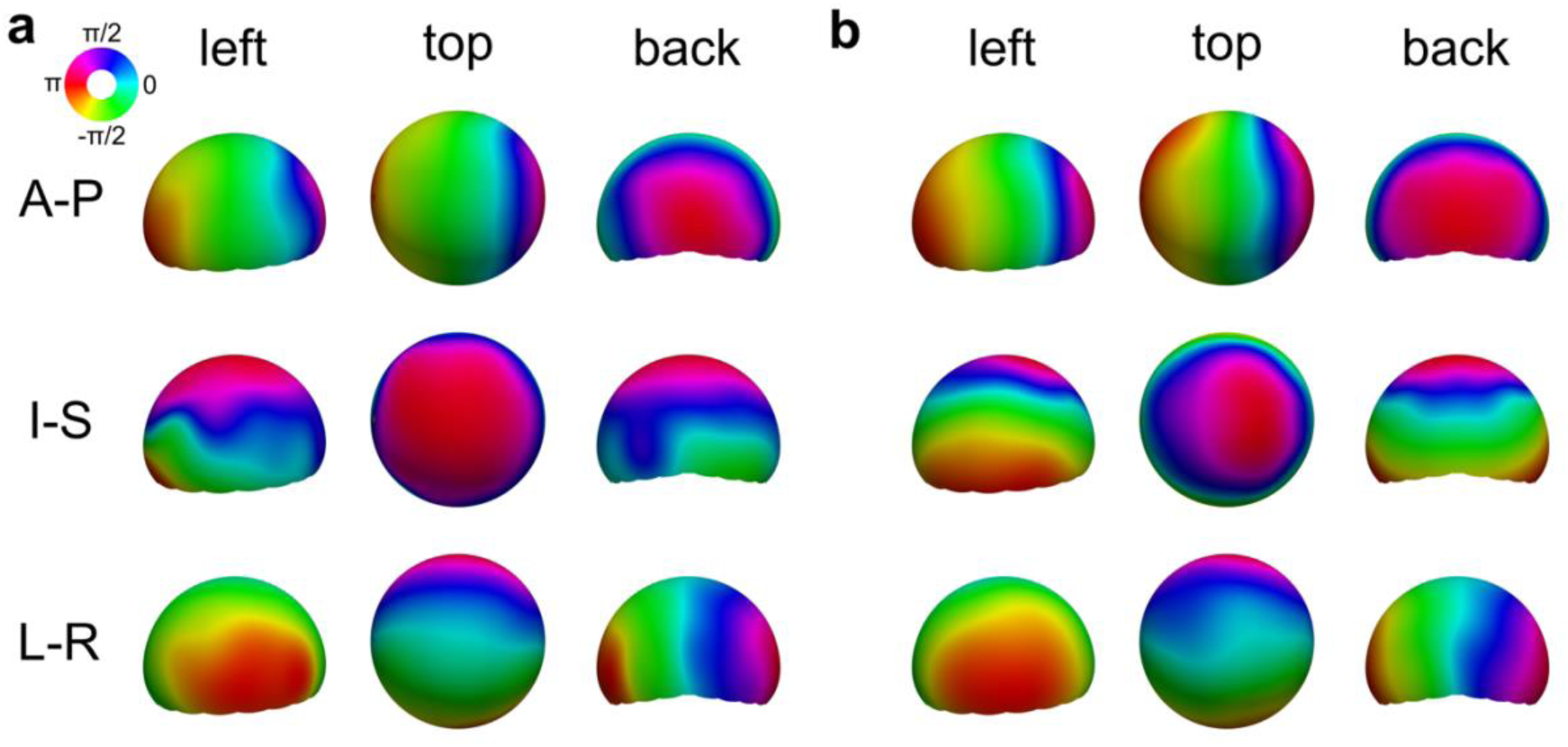
Global traveling waves components for two example participants. **a** Global traveling wave components projected onto cardinal axes for an example participant. A-P: anterior-posterior; I-S: inferior-superior; L-R: left-right. **b** Same as **a**, for another example participant.

**Supplementary Fig. S4.**
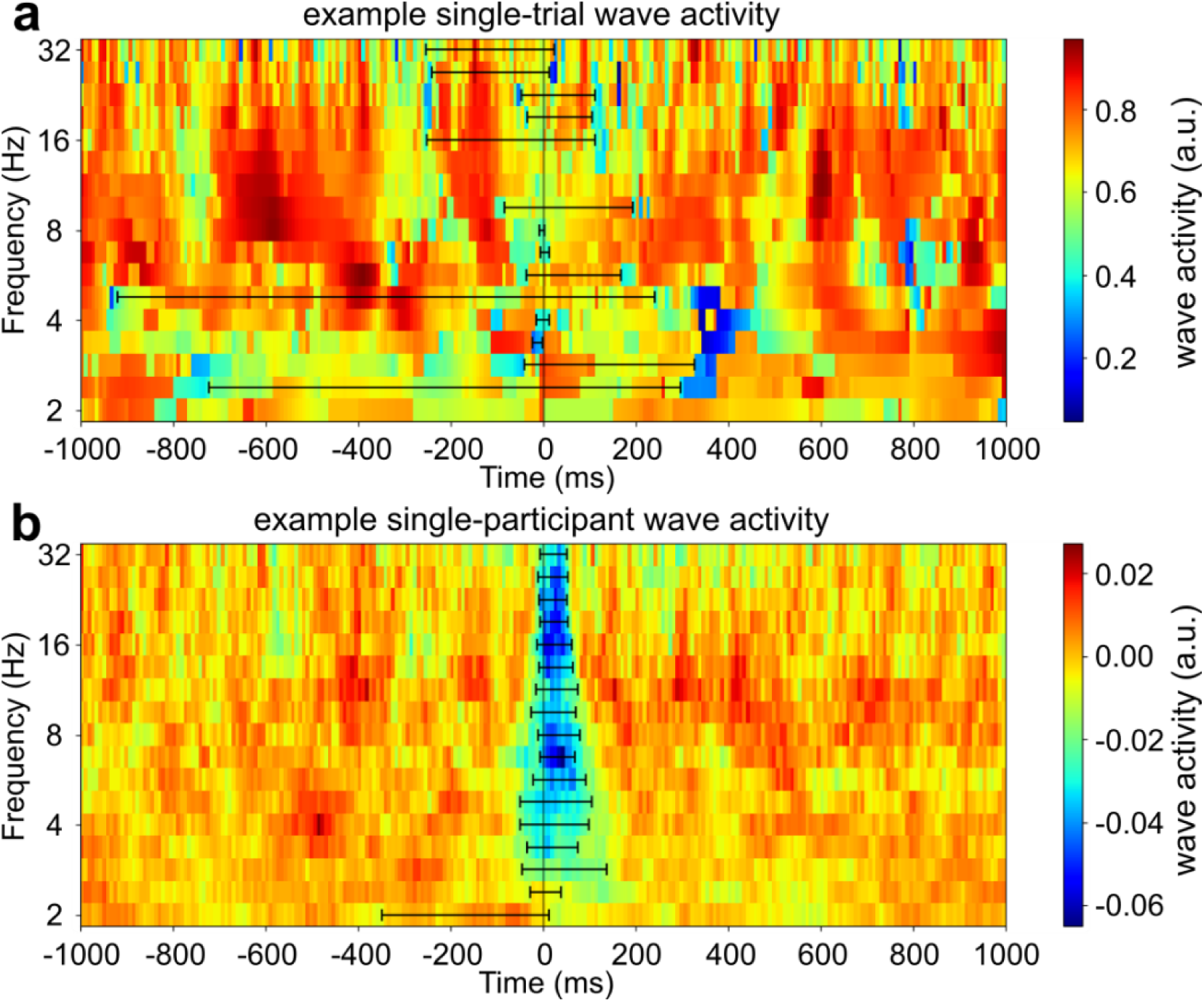
Wave activity is a continuously measured, episodic feature, present throughout the epoch from single trials to averaged trials. **a** Example single-trial wave activity across time and frequency (rFEF-TMS condition). The solid black vertical lines mark TMS pulse onset. Black horizontal bars mark the time-frequency extent of the disruption identified at the single-trial level, for illustration. **b** same as **a**, averaged across trials for an example participant.

**Supplementary Fig. S5.**
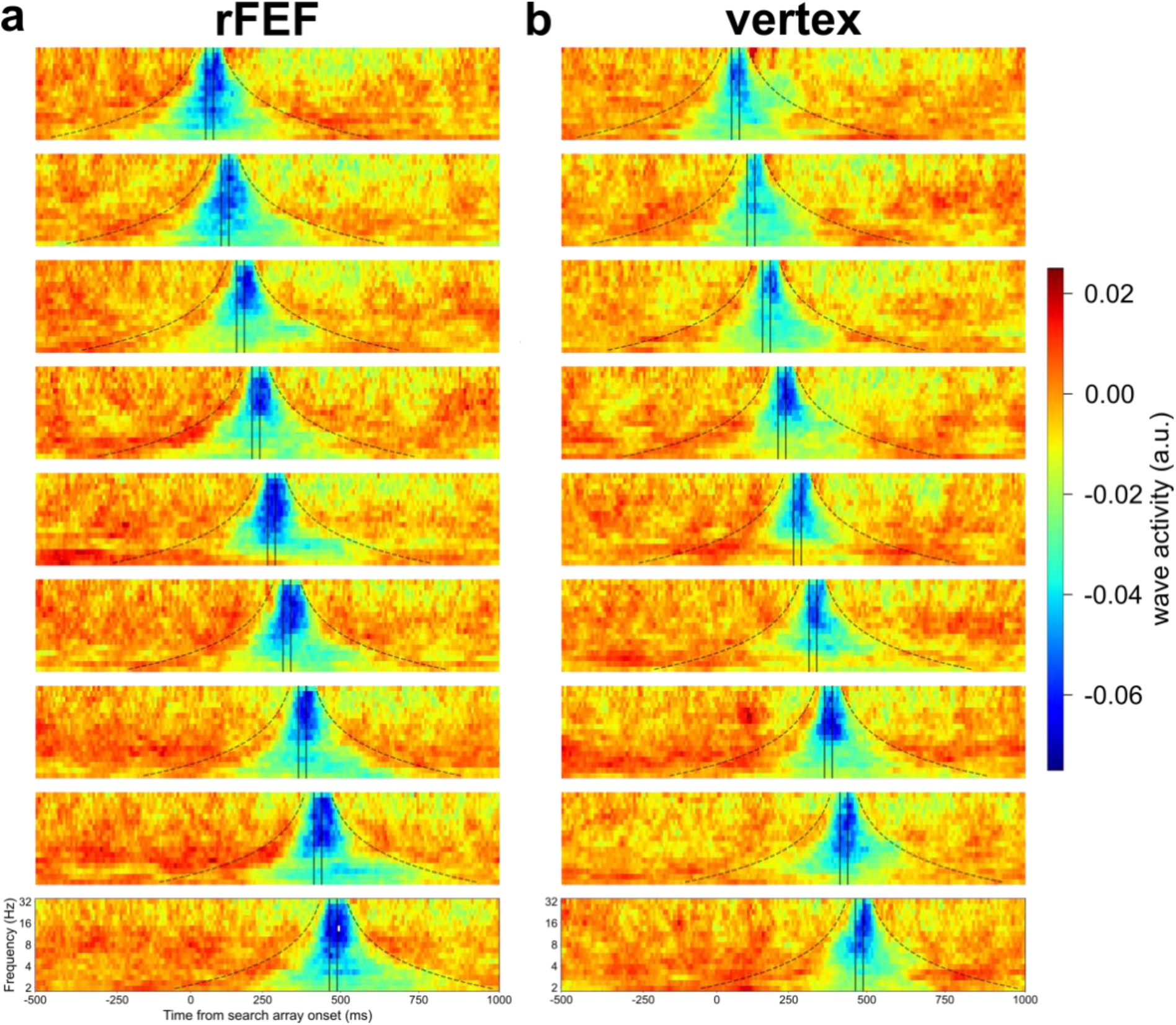
Disruption of global wave activity shifts across TMS latencies. **a** Global traveling wave activity across time and frequency in the rFEF-TMS condition. The solid black lines mark TMS pulse onsets. The cone of influence is overlayed. TMS transiently disrupts global wave activity, and this disruption is temporally specific to the TMS pulses, as evidenced by its systematic temporal shift across different TMS latencies. **b** Same as **a**, but for the Vertex-TMS condition.

**Supplementary Fig. S6.**
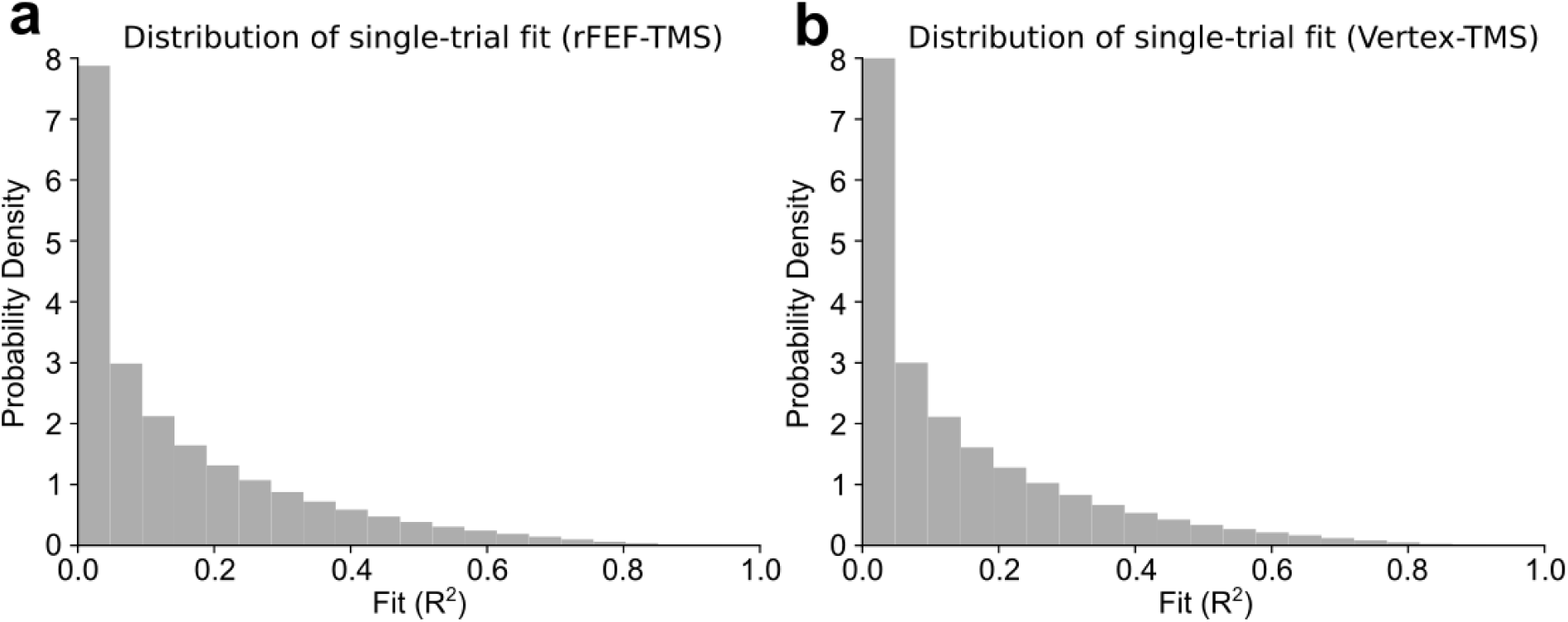
Single-trial anterior-to-posterior wave activity is continuously distributed, not bimodal. Distribution of single-trial anterior-to-posterior wave activity (fit, R^2^) around TMS onset (−300 to 400 ms, 2–32 Hz), pooled across participants, for the rFEF-TMS (**a**) and Vertex-TMS (**b**) conditions. In both conditions, the distribution is continuous and unimodal, with most single-trial values concentrated near zero with a tail toward higher values, rather than showing a bimodal (present/absent) pattern. This indicates that wave activity is a continuously varying measure of spatial phase organization rather than a binary detection of discrete wave events.

**Supplementary Fig. S7.**
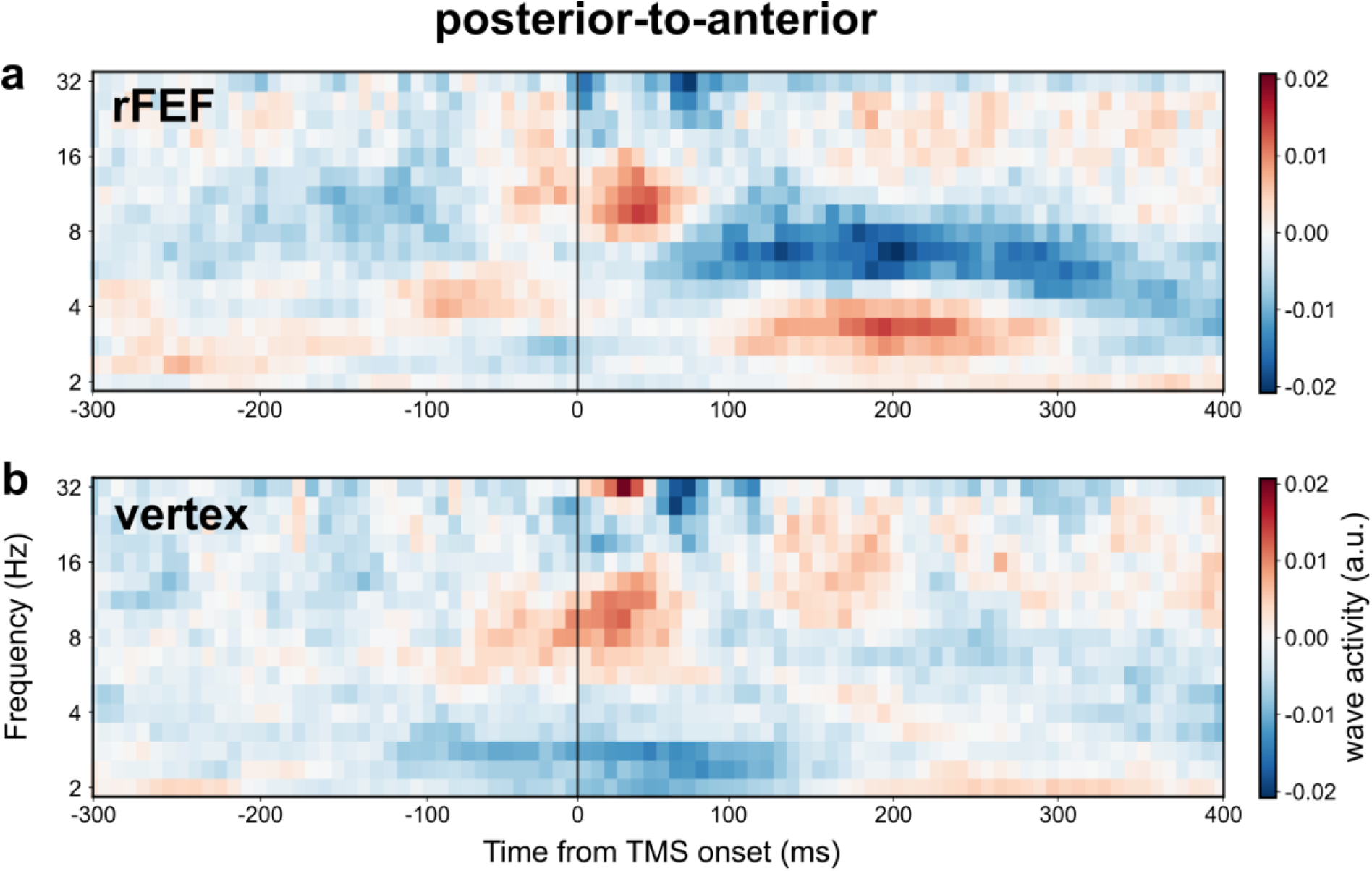
Posterior-to-anterior traveling wave activity following TMS. **a** Posterior-to-anterior traveling wave activity across time and frequency in the rFEF-TMS condition. The solid black line marks TMS onset (first pulse). **b** Same as **a**, but for the Vertex-TMS condition. In contrast to the anterior-to-posterior direction, no significant clusters were observed in the theta band for the posterior-to-anterior direction.

**Supplementary Fig. S8.**
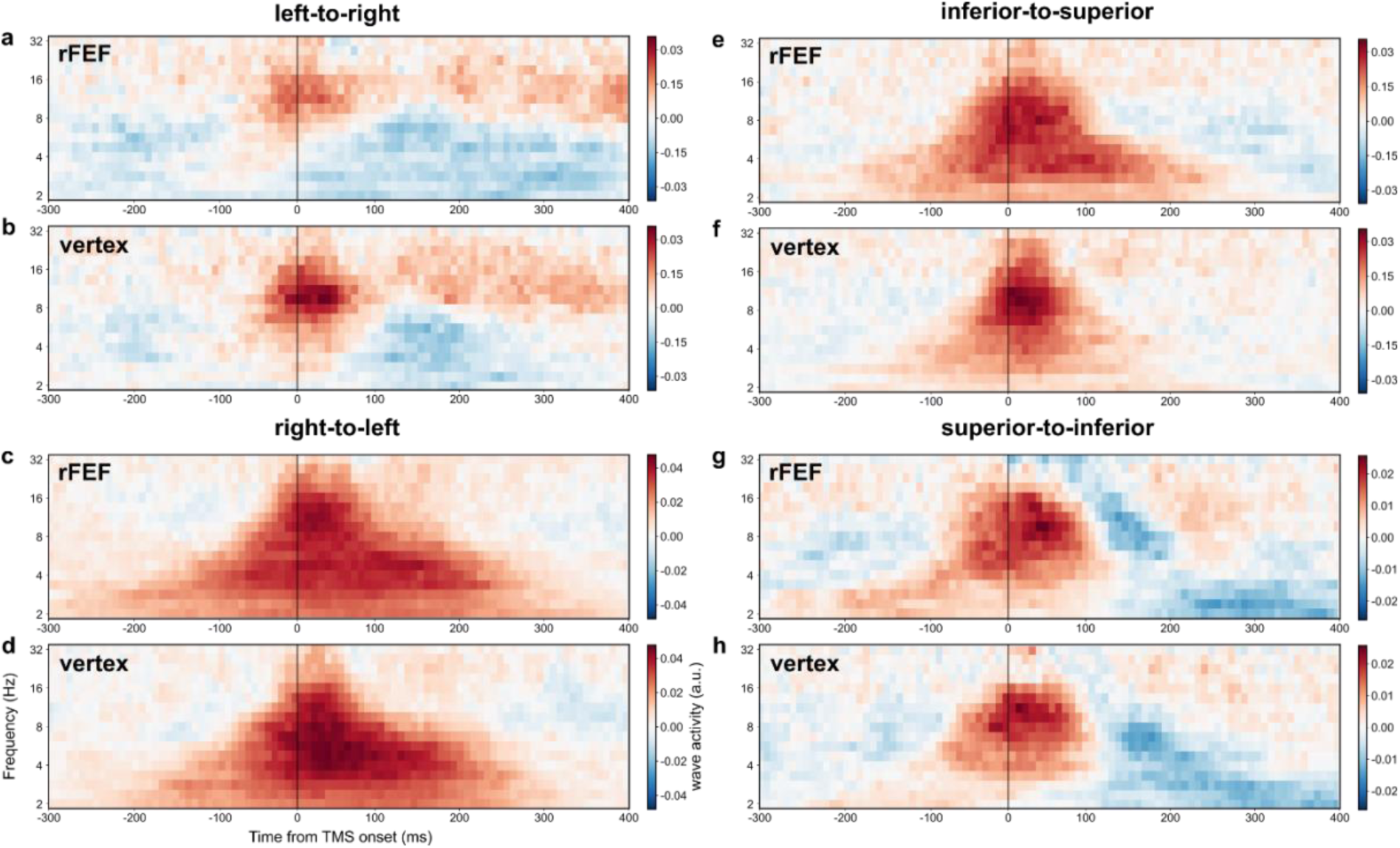
Traveling wave activity along other Cartesian axes following TMS. **a** Left-to-right traveling wave activity across time and frequency in the rFEF-TMS condition. The solid black line marks TMS onset (first pulse). **b** Same as **a**, but for the Vertex-TMS condition. **c** Right-to-left traveling wave activity across time and frequency in the rFEF-TMS condition. **d** Same as **c**, but for the Vertex-TMS condition. **e** Inferior-to-superior traveling wave activity across time and frequency in the rFEF-TMS condition. **f** Same as **e**, but for the Vertex-TMS condition. **g** Superior-to-inferior traveling wave activity across time and frequency in the rFEF-TMS condition. **h** Same as **g**, but for the Vertex-TMS condition. None of these Cartesian directions exhibited significant cluster-level difference in theta traveling waves between rFEF-TMS and Vertex-TMS conditions.

